# *C. elegans* RHY-1 and CYSL-1 act independently of HIF-1 to promote survival in hydrogen sulfide

**DOI:** 10.1101/628784

**Authors:** Joseph W. Horsman, Frazer I. Heinis, Dana L. Miller

## Abstract

Hydrogen sulfide (H_2_S) is an endogenously produced signaling molecule that can be cytoprotective, especially in conditions of ischemia/reperfusion injury. However, exposure to exogenous H_2_S can be toxic, perhaps due to unregulated activation of endogenous H_2_S signaling pathways. We use the nematode *C. elegans* to define mechanisms that mediate the physiological effects of H_2_S in animals. We have previously shown that in *C. elegans* the hypoxia inducible factor (*hif-1*) coordinates the initial transcriptional response to H_2_S and is essential to survive exposure to low concentrations of H_2_S. In this study, we performed a forward genetic screen to identify mutations that suppress the lethality of *hif-1* mutant animals in H_2_S. The mutations we recovered do not suppress embryonic lethality or reproductive arrest of *hif-1* mutant animals in hypoxia, nor can they improve viability of *hif-1* mutant animals exposed to hydrogen cyanide, indicating that these are specific for H_2_S. We found that the *hif-1* suppressor mutations activate the *skn-1*/Nrf2 transcription factor. Activation of SKN-1 by *hif-1* suppressor mutations increased the expression of a subset of H_2_S-responsive genes, consistent with our previous finding that *skn-1* plays a role in the transcriptional response to H_2_S. Using transgenic rescue, we show a single gene, *rhy-1*, alone is sufficient to protect *hif-1* mutant animals in H_2_S. Our data indicate that RHY-1 acts in concert with CYSL-1, an orthologue of human cystathionine β-synthase, to promote survival in H_2_S. The *rhy-1* gene encodes a predicated O-acyltransferase enzyme that has previously been shown to negatively regulate HIF-1 activity. Our studies reveal a novel function of RHY-1, which is independent of *hif-1*, that protects against toxic effects of H_2_S.

## Introduction

Hydrogen sulfide (H_2_S) is endogenously produced by animal cells that has various roles in cellular signaling (Li et al., 2011). Exposure to low levels of exogenous H_2_S induces a suspended-animation like state in rodents that is associated with increased survival in whole-body hypoxia and improved outcome in models of ischemia-reperfusion injury (Blackstone and Roth, 2007; Jensen et al., 2017; Wu et al., 2015). However, H_2_S is also a common environmental toxin, and human exposure is associated with respiratory and neurological dysfunction (Malone Rubright et al., 2017). In *C. elegans*, exogenous H_2_S increases lifespan and protects from hypoxia-induced protein aggregation (Miller and Roth, 2007; Fawcett et al., 2015), suggesting that protective physiological effects of exposure to low H_2_S may be conserved.

In *C. elegans*, the *hif-1* and *skn-1* transcription factors mediate the initial transcriptional response to exogenous H_2_S (Miller et al., 2011). HIF-1 is a conserved bHLH transcription factor that coordinates the transcriptional response to hypoxia in metazoans (Semenza, 2012). All of the genes induced within one hour of exposure to H_2_S are dependent on *hif-1* (Miller et al., 2011), and animals with mutations that disrupt *hif-1* die when exposed to low H_2_S (Budde and Roth, 2010). HIF-1 activity is regulated post-translationally in response to O_2_ availability. HIF-1 is hydroxylated at a specific proline residue by the EGL-9 prolyl hydroxylase in a reaction that requires molecular O_2_, and hydroxylated HIF-1 is recognized by the E3 ubiquitin ligase VHL-1 and subsequently degraded (Epstein et al., 2001; Maxwell et al., 1999). In H_2_S, EGL-9 associates with CYSL-1, one of four *C. elegans* orthologues of human cystathionine β-synthase, and HIF-1 protein accumulates (Budde and Roth, 2010; Ma et al., 2012).

SKN-1 is an orthologue of Nrf2 (Walker et al., 2000) that is required for specification of the pharynx and intestine during embryogenesis (Bowerman et al., 1992). In adults, *skn-1* coordinates a Phase II-like response to oxidative stress (An and Blackwell, 2003). A subset of early transcriptional responses to H_2_S are disrupted by depletion of *skn-1*, and *skn-1* mutant animals die when exposed to low H_2_S (Miller et al., 2011). Postembryonic SKN-1 activity is regulated by phosphorylation (An et al., 2005; Kell et al., 2007) and association with the WD40 repeat protein, WDR-23, which promotes the degradation of SKN-1 (Choe et al., 2009). In mammals, Nrf2 is activated by H_2_S and is associated with protective effects of H_2_S in myocardial ischemia and oxidative stress resistance (Calvert et al., 2009; Yang et al., 2013).

In this study, we used a forward genetic screen to find mutations that suppress the lethality of *hif-1* mutant animals in low H_2_S. Although the entire transcriptional response to H_2_S is abrogated in *hif-1* mutant animals (Miller et al., 2011), we recovered six mutations that by-pass the requirement for *hif-1* in H_2_S. Cloning these mutations identified two gain-of-function alleles of *skn-1* and three loss-of-function alleles of *wdr-23*. The mutations we recovered are specific for the function of HIF-1 in H_2_S, as they do not suppress *hif-1* phenotypes in hypoxia or cyanide. We demonstrate that the protective effects of increased SKN-1 activity are due to increased expression of *rhy-1*. The *rhy-1* gene encodes a predicted acyltransferase that functions as a negative regulator of HIF-1 (Shen et al., 2006). We show that expression of *rhy-1* is both necessary and sufficient for *hif-1(ia04)* mutant animals to survive exposure to H_2_S. RHY-1 requires CYSL-1, a cysteine synthase-like factor, but not the EGL-9 prolyl hydroxylase, to suppress the lethality of *hif-1(ia04)* mutant animals expose to H_2_S. Together, our data demonstrate that RHY-1 has a previously unidentified function that promotes survival in H_2_S, independently of HIF-1.

## Results

### Activation of SKN-1 suppresses lethality of *hif-1(ia04)* in H_2_S

In *C. elegans*, the conserved transcription factor HIF-1 is essential to survive exposure to low H_2_S (Budde and Roth, 2010). In order to identify genetic factors that work with HIF-1 in H_2_S, we performed a forward-genetic screen to recover mutations that suppressed the lethality of *hif-1(ia04)* mutant animals exposed to 50 ppm H_2_S. The *hif-1(ia04)* mutation is a predicted null allele, removing 1,231 bp including 3 exons and is a predicted molecular null (Jiang et al., 2001). We chose to use 50 ppm H_2_S, as this concentration is invariably lethal to *hif-1* animals but increases lifespan and reduces protein aggregation in wild-type animals (Fawcett et al., 2015; Miller and Roth, 2007). We hypothesized that genes identified by this screen would act either downstream or in parallel to HIF-1 in the response to H_2_S.

Our screen isolated 6 independent alleles that bypassed the requirement for HIF-1 in H_2_S. We refer to these as *suh* mutations, for <u>su</u>ppressor of *<u>h</u>if-1* in H_2_S (Figure 1A). In the screen for the *suh* alleles, we isolated 15 surviving animals after exposing 620,000 mutagenized P0 *hif-1(ia04)* to H_2_S. Ten of these mutants had the same genetic lesion suggesting a “jackpot”, where multiple animals contain a lesion from a single mutagenic event. We included only one representative of this group (*uwa02*) in our studies. The *suh* mutations do not rescue survival of embryos in H_2_S, as *hif-1(ia04); suh* mutant embryos laid in the H_2_S environment were not viable (data not shown).

**Figure 1.**
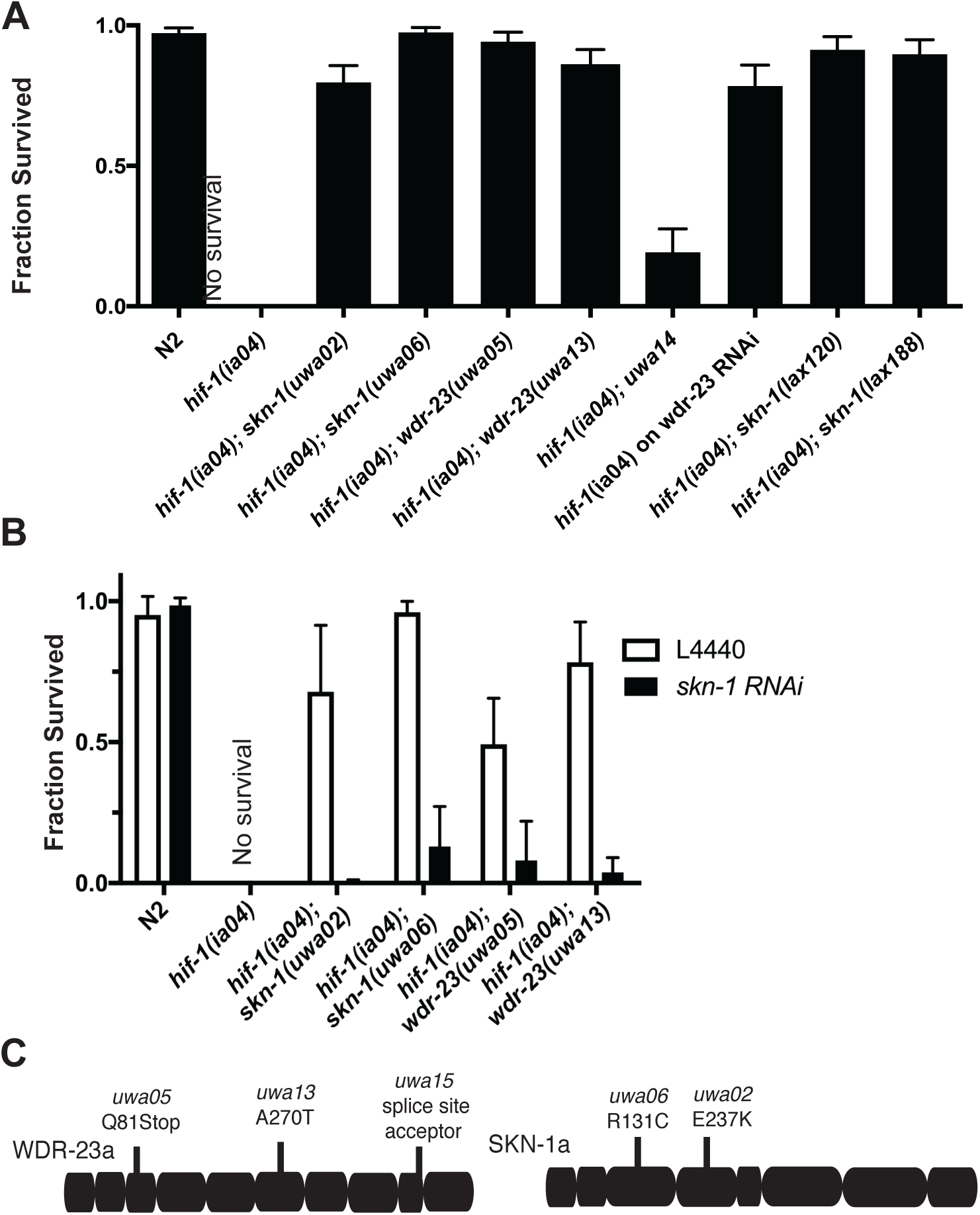
Screen to suppress *hif-*1 lethality in H_2_S isolated mutations that increase SKN-1 activity. **A.** Mutant strains isolated in a forward genetic screen for <u>su</u>ppressors of *<u>h</u>if-1* lethality in H_2_S (Suh phenotype). The mean survival of recovered mutant strains is shown after 16 h in 50 ppm H_2_S. N2 is wildtype, *lax* alleles are *skn-1* gain-of-function alleles from other groups (Paek et al., 2012). Data are average of 10-13 independent experiments, and each replicate experiment included at least 10 animals. Error bars are SEM. **B.** *skn-1* is required for *suh* mutant survival in H_2_S. Animals were grown on *skn-1(RNAi)* food (filled bars) or control L4440 food (open bars) and then exposed to 50 ppm H_2_S overnight. Data are average of at least 3 independent experiments. Error bars are standard deviation of the mean. **C.** Mutations in *wdr-23* and *skn-1* that confer Suh phenotype. Schematic shows exon structure of *wdr-23* and *skn-1* (www.wormbase.org WS204). Alleles isolated in this screen are indicated above the schematic. Recovered loss-of-function mutations in *wdr-23* (*uwa05, uwa13*, and *uwa15*) would affect all 9 predicted isoforms. Gain of function mutations in *skn-1* (*uwa02* and *uwa06*) would affect a and c isoforms only.

All of the recovered alleles were recessive for suppressing loss of *hif-1* in H_2_S except *uwa06*. The five recessive mutations were grouped into three complementation groups by measuring the survival of trans-heterozygotes of different Suh alleles (Supplemental Figure 1). One group included *uwa05, uwa15* and *uwa13*, which all failed to complement one another. The other two complementation groups included one allele each, *uwa14* and *uwa02*. These data indicate our screen is not saturation, and suggest there could be other genes that could mutate to the Suh phenotype.

We cloned the mutations that confer the Suh phenotype using whole genome sequencing, comparing mutant sequences (*uwa02, uwa14, uwa05* and *uwa15*) to the parental *hif-1(ia04)* strain (Minevich et al., 2012). We found that *uwa05* and *uwa15*, which are in the same complementation group, both contained variations in the *wdr-23* gene: in *uwa05* we found a C to T transition that changed Q81 to an ochre stop, and in *uwa15* strain there was a G to A transition that disrupts a splice-site donor between the 9^th^ and 10^th^ exon of *wdr-23* (Figure 1C). We directly sequenced the *wdr-23* locus of *uwa13*, the third mutant strain in this complementation group, and identified a G to A transition that introduced an A270T missense mutation in the seventh exon, near several other missense mutations that have been shown to disrupt WDR-23 function (Figure 1C; Hasegawa and Miwa, 2010). Because these mutations were all recessive and included one allele with a premature stop codon, we predicted they would be loss-of-function alleles of *wdr-23*. To test this possibility, we asked whether *wdr-23(RNAi)* would recapitulate the Suh phenotype in *hif-1(ia04)* mutant animals. Consistent with this hypothesis, we found that 78% of *hif-1(ia04)* animals grown on *wdr-23* RNAi survived exposure to 50 ppm H_2_S, whereas none of the animals grown on the control RNAi survived (Figure 1A). We conclude that disruption of *wdr-23* bypasses the requirement for HIF-1 in H_2_S.

WDR-23 is a WD-40 repeat protein that acts with the CUL-4/DDB-1 ubiquitin ligase to target SKN-1 for degradation, acting analogous to Keap1 regulation of Nrf2 in mammals (Choe et al., 2009; Kobayashi et al., 2004). Reduction in WDR-23 function leads to an increase in SKN-1 transcriptional activity and increased stress resistance, and lifespan, and these *wdr-23*-knockout phenotypes are dependent on *skn-1* (Choe et al., 2009; Tang and Choe, 2015). We had previously found that *skn-1* was involved in mediating the transcriptional response to H_2_S (Miller et al., 2011). These facts suggested that increased SKN-1 activity suppressed the lethality of *hif-1(ia04)* mutant animals in H_2_S. Consistent with this hypothesis when we depleted *skn-1* by RNAi the Suh phenotype of *uwa06, uwa02*, and *wdr-23* was abrogated (Figure 1B). We also asked if expression of the P*gst-4::gfp* fluorescent reporter, which exhibits increased fluorescence in conditions where SKN-1 activity is increased (An and Blackwell, 2003), was induced in animals with *suh* alleles (Supplemental Figure 2). As expected from our previous microarray data, P*gst-4::gfp* is not induced by exposure to 50 ppm H_2_S in wild-type animals (Miller et al., 2011). However, we observed strong expression of the P*gst-4::gfp* reporter in Suh strains without exposure to H_2_S. In *wdr-23* mutant animals (*suh* alleles *uwa05, uwa13* and *uwa15*), GFP expression was only observed in homozygous mutant animals, whereas the *skn-1(gf)* alleles *uwa06* and *uwa02* increased expression of the reporter even as heterozygotes.

We reasoned that if increased SKN-1 activity could suppress the requirement for HIF-1 in H_2_S, some of the *suh* alleles we recovered may be gain-of-function mutations in SKN-1. Previous screens for mutations that increase expression of *Pgst-4::gfp* had identified gain-of-function mutations in SKN-1 (Paek et al., 2012). Consistent with this, we found a G at A transition in the *skn-1* coding sequence that introduced an E237K substitution in the whole-genome sequence of the *uwa02* strain and, by directly sequencing the *skn-1* locus, we found the dominant *uwa06* allele contains an R131C substitution in *skn-1* (Figure 1C). The E237K substitution in *uwa02* is the same change as reported for *skn-1(lax188*) gain-of-function allele (Paek et al., 2012). The fact that expression of *Pgst-4::gfp* is expressed in animals with the *uwa02* and *uwa06* alleles (Supplemental Figure 2) is consistent with previous observations that *skn-1(gf)* alleles can act dominantly to increase transcription of targets (Paek et al., 2012). We also determined that previously isolated *skn-1(gf)* alleles, *lax188* and *lax120*, both suppressed the lethality of *hif-1(ia04)* animals in H_2_S (Figure 1A). We conclude that activation of SKN-1 promotes survival in H_2_S independently of HIF-1.

Despite whole genome sequencing, *uwa14*, remains uncloned. We did not find any changes in either the *wdr-23* or *skn-1* loci in this strain, and this allele does not induce expression of *Pgst-4::gfp*. These data suggest that *uwa14* acts in a separate pathway from *wdr-23* and *skn-1*. We have not further characterized this allele at this time.

### Suppressors of *hif-1* lethality specifically promote survival in H_2_S

Although we have focused on the role of HIF-1 in H_2_S, the conserved HIF-1 transcription factor is best known for its role in coordinating the transcriptional response to hypoxia. In *C. elegans, hif-1* is required for embryos to survive exposure to hypoxia (Jiang et al., 2001; Nystul and Roth, 2004). We found that *hif-1(ia04); suh* mutant embryos died in hypoxia similar to *hif-1(ia04)* embryos (Figure 2A). This result suggests that activation of SKN-1 is not sufficient to compensate for the loss of *hif-1* in hypoxia. Alternatively, it could be that this mechanism is not active during embryogenesis. *skn-1* is required during embryogenesis for the appropriate development of the mesendoderm, which may preclude its activation in response to stress (Bowerman et al., 1992). To address this possibility, we measured egg-laying of adult *C. elegans* in 5,000 ppm O_2_. In this condition, wild-type animals continue to produce and lay eggs, while *hif-1(ia04)* animals enter into a hypoxia-induced diapause (Miller and Roth, 2009). We found egg-laying in *hif-1(ia04); suh* double mutants was similar to the *hif-1(ia04)* animals (Figure 2B). Together, these data are consistent with the interpretation that *suh* alleles do not suppress the loss of *hif-1* in hypoxia. The fact that activating SKN-1 suppresses the loss of *hif-1* in H_2_S but not hypoxia supports the hypothesis that different stresses result in activation of distinct HIF-1-mediated responses. This is consistent with our previous observation that there is little overlap in the targets of the *hif-1*-dependent transcriptional responses to hypoxia and H_2_S (Miller et al., 2011).

**Figure 2.**
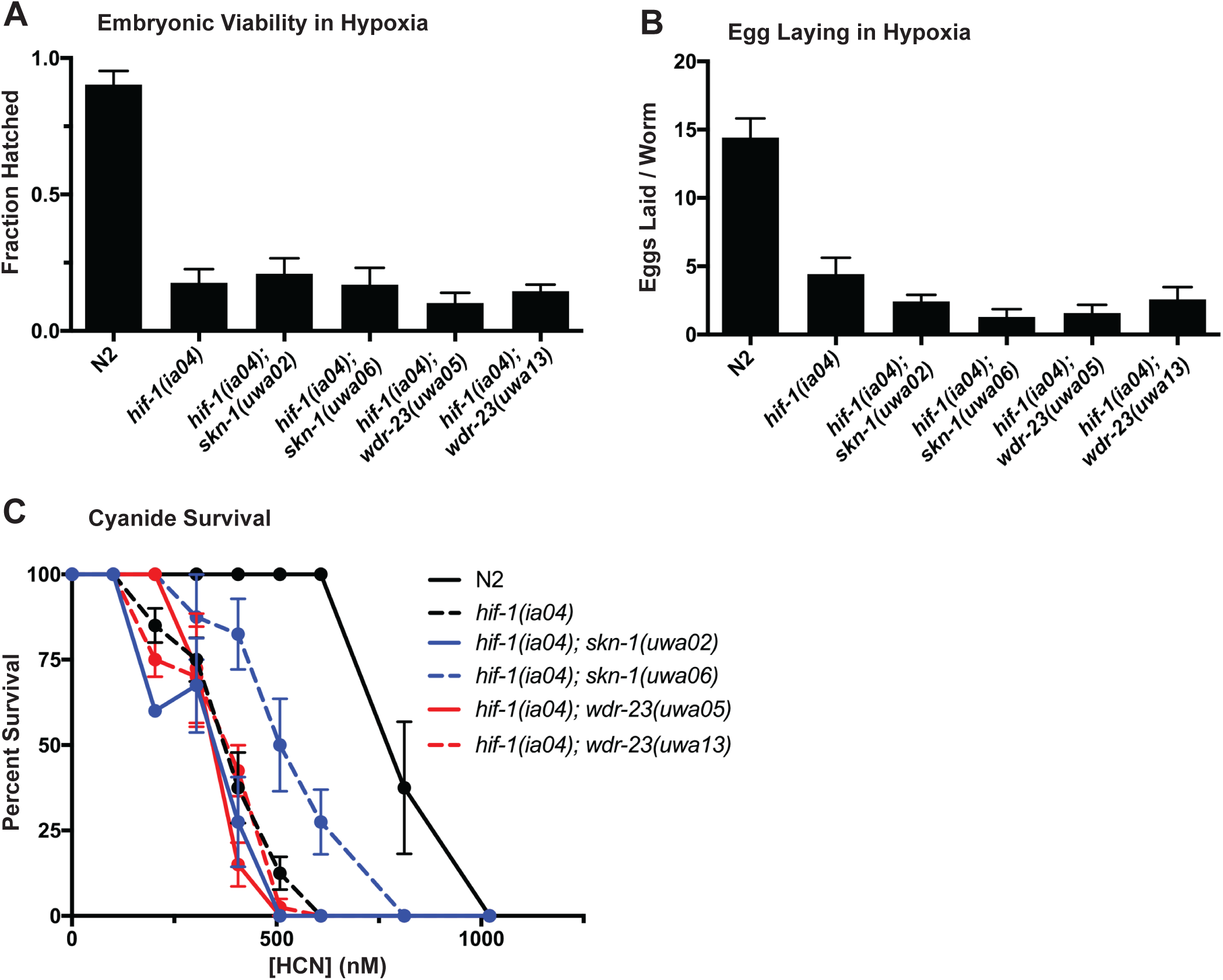
Mutations in *wdr-23* and *skn-1* specifically suppress H_2_S phenotypes of *hif-1(ia04)* mutant animals. **A.** Embryonic lethality of hif-1(ia04) mutant animals exposed to hypoxia is not rescued by *suh* alleles. Embryos on plates were exposed to 5,000 ppm O_2_ for 24 h, and the fraction that hatched was scored upon removal from hypoxia. Each data set includes at least 10 independent experiments, except *hif-1(ia04); wdr-23(uwa05)*, which includes 6 replicates. Graph show mean viability, error bars are SEM. **B.** Hypoxia-induced arrest of egg-laying is not suppressed by *suh* alleles. First day adults were exposed to 5,000 ppm O2 for 20h, and the number of eggs laid was scored. Data are average of 7 independent replicates, with 6 animals for each strain per replicate. Error bars show SEM. **C.** *suh* alleles do not suppress *hif-1(ia04)* sensitivity to HCN. Animals were exposed to each concentration of HCN overnight, and then the number of animals surviving was scored. Each data point is the mean of 2 (0 nM and 100 nM) or 4 (all other HCN concentrations) independent replicates, error bars are SEM.

In addition to its role in mediating the transcriptional response to hypoxia, HIF-1 mediates the response to hydrogen cyanide (HCN) and protects against fast-killing by *Pseudomonas aeruginosa* (Darby et al., 1999; Gallagher and Manoil, 2001; Shao et al., 2010). HCN has been proposed to share the same mechanisms of toxicity as H_2_S (Cooper and Brown, 2008), and genetic evidence suggests that *C. elegans* have a common response to H_2_S and HCN (Budde and Roth, 2011). In this scenario, we would predict that the *suh* alleles would also confer protection against HCN. However, when we measured the survival of animals exposed to gaseous HCN we observed that most *hif-1(ia04); suh* double mutants were equally sensitive to HCN as *hif-1(ia04)* single mutant animals (Figure 2C). The one exception was the *hif-1(ia04); skn-1(uwa06)* double mutant, which has a slight increase in viability relative to *hif-1(ia04)* animals. These data show that the *hif-1*-mediated resistance to H_2_S and HCN toxicity are genetically separable, which argues against the hypothesis that there is a common mechanism for detoxification of HCN and H_2_S. This observation is consistent with the fact that treatments for cyanide poisoning are not generally effective at treating and H_2_S toxicity in mammals (Jiang et al., 2016).

### *rhy-1* is necessary and sufficient to suppress *hif-1* lethality in H_2_S

Both HIF-1 and SKN-1 are transcription factors that mediate early transcriptional responses to H_2_S (Miller et al., 2011), suggesting the possibility that activation of SKN-1 by the *suh* alleles could induce expression of H_2_S-responsive genes required for survival. We first asked if any of the known H_2_S-inducible genes were constitutively induced by activation of *skn-1* (Figure 3A). We measured transcript abundance of H_2_S-induced genes in *hif-1(ia04); skn-1(uwa02)* double mutant animals using qRT-PCR (Miller et al., 2011). In animals that had not been exposed to H_2_S, four of these transcripts were more abundant in *hif-1(ia04); skn*-*1(uwa02)* animals compared to N2 controls: *nspe-3, nit-1, dhs-8* and *rhy-1*. To determine if expression of these genes alone is sufficient to confer the Suh phenotype, we generated extrachromosomal arrays expressing all four of the genes up-regulated in the *skn-1(uwa02)* mutation (Figure 3B). We also included R08E5.1 in these arrays based on preliminary experiments that suggested it was induced by H_2_S in *hif-1(ia04); skn-1(uwa02)* mutant animals (this result was not reproducible). Each gene, with its native promoter and 3’ UTR, was amplified from genomic DNA and the PCR products mixed and injected into the gonads of *hif-1(ia04)* animals to generate a multi-copy extrachromosomal array, *uwaEx09*, that drove expression of these five genes. The *hif-1(ia04); uwaEx09* animals survived exposure to H_2_S (Fig 3B), suggesting that increased expression of one or more of the genes induced by activation of SKN-1 accounts for the observed Suh phenotype.

**Figure 3.**
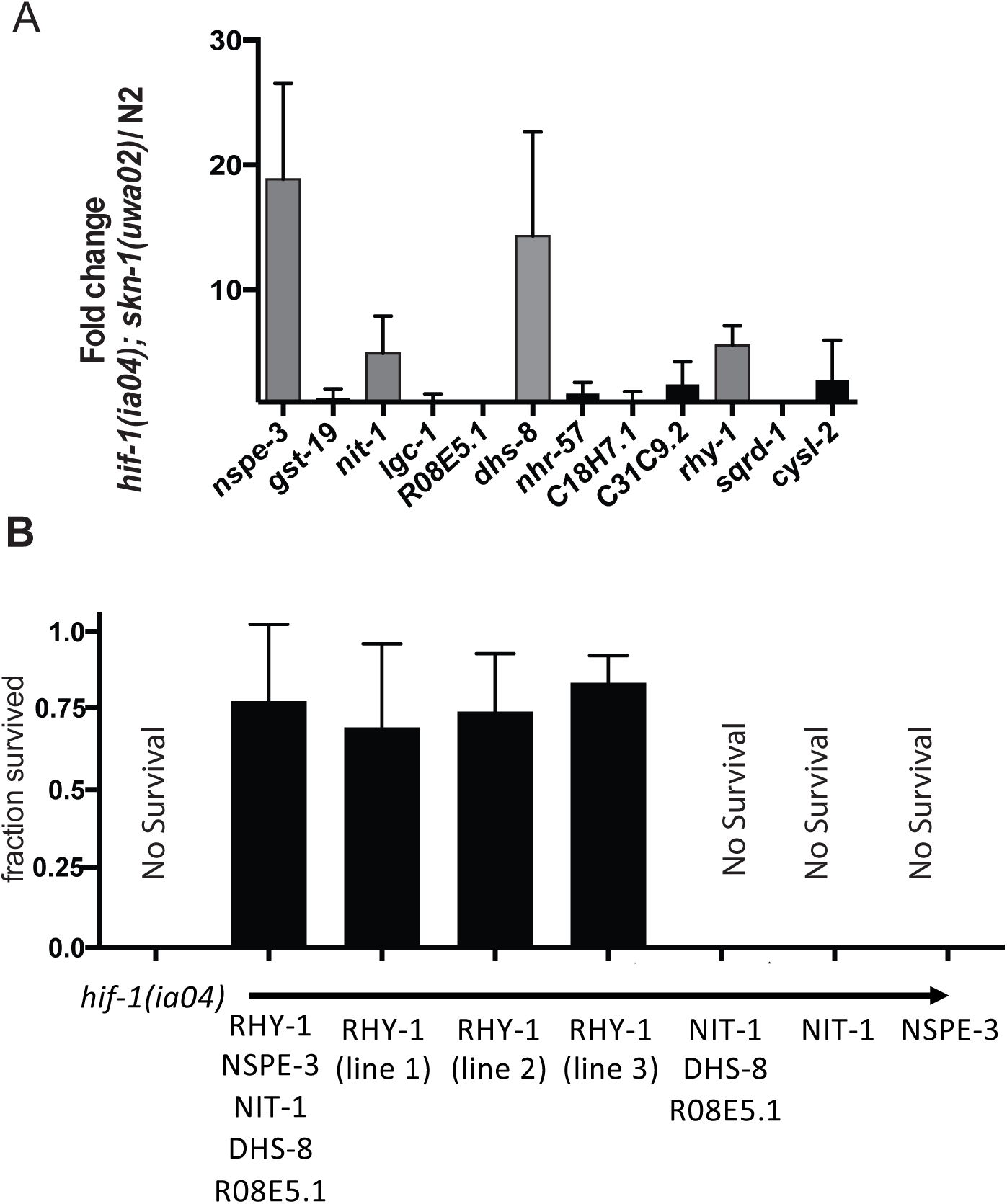
*rhy-1* expression is sufficient to suppress *hif-1* lethality in H_2_S. A. H_2_S-induced genes are constitutively expressed in *skn-1(uwa02gf)* mutant animals. The mRNA abundance of each gene was measured by qRT-PCR in *hif-1(ia04); skn-1(uwa02)* and N2. Each measurement was repeated in three independent experiments with three technical replicates in each experiment. Relative expression levels were calculated with ΔΔC_t_, graph shows average fold change between strains. Error bars are SD. **B.** Transgenic expression of RHY-1 suppresses *hif-1* lethality in H_2_S. Genomic DNA for each gene was amplified and injected into *hif-1(ia04)* mutant animals to generate multi-copy extrachromosomal arrays. Cohorts of animals with each array were exposed to H_2_S overnight as L4. Graph shows average survival from 4-8 independent experiments, with at least 10 animals in each experimental test. Error bars are SEM. All extrachromosomal arrays were in the *hif-1(ia04)* mutant background; genes included in each array are indicated below the graph.

To further define which of the transcripts up-regulated by activation of SKN-1 are required for *hif-1* mutant animals to survive exposure to H_2_S, we generated transgenic animals with different combinations of the 5 genes in the original array (Figure 3B). We found that two separate arrays containing only genomic *rhy-1 (uwaEx10* and *uwaEx11)* rescued the lethality of *hif-1(ia04)* mutant animals in H_2_S (Figure 3B). All other arrays containing either a single genes or combinations of genes that did not include *rhy-1* did not suppress lethality of *hif(ia04)* animals in H_2_S (Figure 3B). This result indicates increased expression of *rhy-1* alone is sufficient to bypasses the requirement for *hif-1* in H_2_S. Consistent with this assertion, we found that *rhy-*1 mRNA was more abundant in the *wdr-23(uwa05), skn-1(uwa06)* and *wdr-23(uwa13)* Suh strains (Supplemental Figure 4). To determine if *rhy-1* was also necessary for the Suh phenotype, we exposed *hif-1(ia04); rhy-1(ok1402)* mutant animals to *wdr-23(RNAi)* to increase the activity of SKN-1. We found that *hif-1(ia04); rhy-1(ok1402)* double mutant animals on *wdr-23* RNAi died when exposed to H_2_S, whereas *hif-1(ia04)* single mutants in the same experiment survived as expected (Figure 4A). Together, these results show that *rhy-1* is both necessary and sufficient to bypass the requirement of *hif-1* in H_2_S.

**Figure 4.**
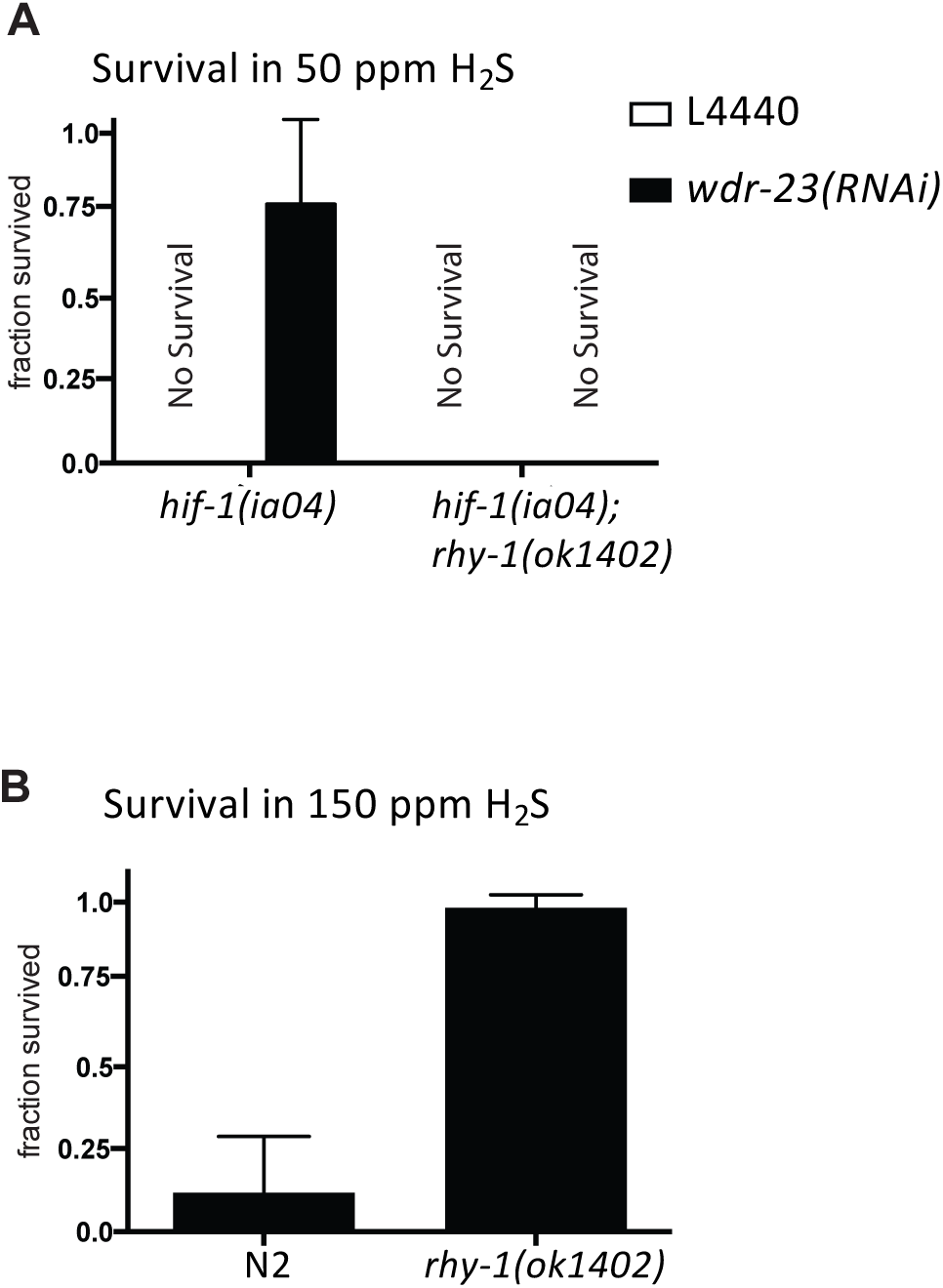
*rhy-1* is necessary to suppress loss of *hif-1* by increased *skn-1*. A. RNAi of *wdr-23* requires *rhy-1* to suppress *hif-1* lethality in H_2_S. Cohorts of *hif-1(ia04)* and *hif-1(ia04)*; *rhy-1(ok1402)* mutant animals were grown on *wdr-23* RNAi food or L4440 control food and then exposed to 50 ppm H_2_S overnight as L4/young adult. The number of animals that survived exposure was scored. Data are from 3 (L4440 controls) or at least 10 (*wdr-23* RNAi experiments) independent replicates, with at least 10 animals in each cohort. Error bars are SEM. **B.** In animals with functional HIF-1, mutations in *rhy-1* promotes survival in high H_2_S. N2 (wild-type) and *rhy-1(ok1402)* mutant animals were exposed to 150 ppm H_2_S overnight, and the number of surviving animals was scored. Data are mean of at least 10 independent experiments with at least 10 animals in each cohort. Error bars show SEM.

### RHY-1 acts with CYSL-1 to promote survival in H_2_S independently of HIF-1

Our data show that RHY-1 is necessary and sufficient *for hif-1(ia04)* mutant animals to survive exposure to H_2_S. This is a new function for RHY-1, which was first characterized as a negative regulator of HIF-1 activity (Ma et al., 2012; Shen et al., 2006). Insofar as constitutive activation of HIF-1 increases survival of *C. elegans* in H_2_S (Budde and Roth, 2010), we predicted that these mutants would be resistant to high H_2_S. Consistent with this, we found that *rhy-1(ok1402)* mutant animals (with wild-type *hif-1*) survive exposure to high H_2_S (150 ppm; Figure 4B). The resistance to high H_2_S depended entirely on *hif-1*, as *hif-1(ia04); rhy-1(ok1402)* double mutants died when exposed to H_2_S. This is similar to increased resistance to H_2_S observed in *egl-9* and *vhl-1* mutant animals (Budde and Roth, 2010), consistent with the notion that RHY-1 is a negative regulator of HIF-1 (Shen et al., 2006).

One mechanism that RHY-1 could be improving survival of *hif-1(ia04)* mutant animals in H_2_S is by increasing the activity of SQRD-1. SQRD-1 is the C. elegans orthologue of the sulfide-quinone oxidoreductase that catalyzes the initial step in the mitochondrial oxidation of sulfide (Hildebrandt and Grieshaber, 2008). In *C. elegans, sqrd-1* upregulation in response to H_2_S exposure requires *hif-1* and sqrd-1 mutant animals die when exposed to H_2_S (Miller and Roth, 2011; Budde and Roth 2011). We reasoned that if RHY-1 was acting through SQRD-1, then *sqrd-1* would be required for the Suh phenotype of *wdr-23(RNAi)*. However, we found that RNAi of *wdr-23* rescued the lethality of *sqrd-1(tm3378)* mutant animals exposed to H_2_S (Figure 5A). Moreover, we did not observe increased abundance of *sqrd-1* transcript in *hif-1(ia04); skn-1(uwa02)* double mutant animals (Figure 3A). These data argue that the survival of *suh* mutant animals in H_2_S does not result from increased detoxification of H_2_S by SQRD-1.

**Figure 5.**
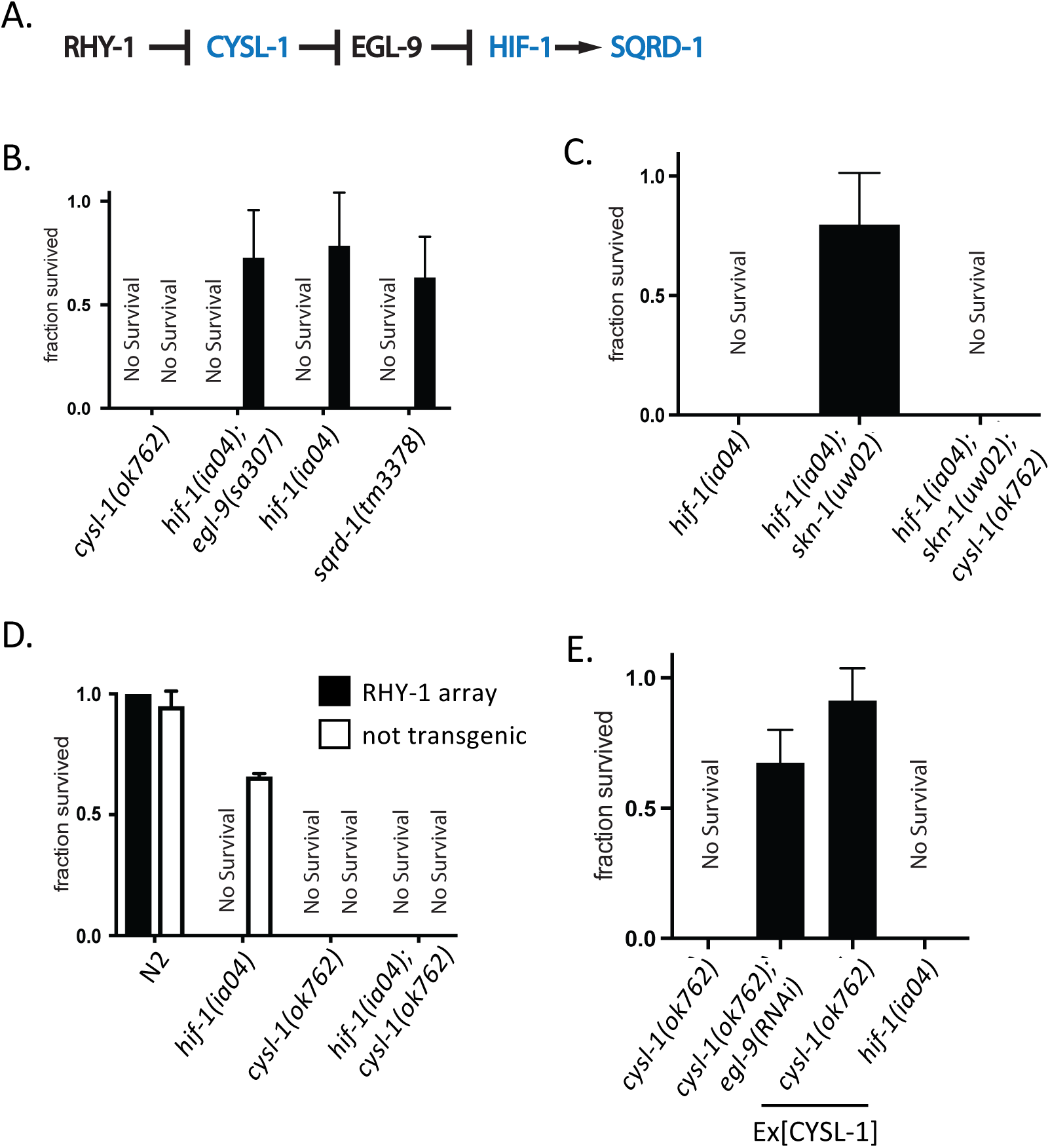
RHY-1 acts with CYSL-1 to promote *hif-1* independent survival in H_2_S. A. Proposed genetic pathway for RHY-1 regulation of HIF-1, adapted from (Ma et al., 2012). Animals with mutations in genes encoding proteins in blue die when exposed to low H_2_S. **B.** Activation of *skn-1* by *wdr-23(RNAi)* requires *cysl-1*, but not *egl-9*, to suppress *hif-1* lethality in H_2_S. Animals of each genotype were grown on *wdr-23(RNAi)* food (filled bars) or L4440 control RNAi food (empty bars) and then exposed to 50 ppm H_2_S overnight. The number of animals that survived exposure was scored. The graph shows mean survival from 3-12 independent replicates with at least 10 animals in each cohort. Error bars are SEM. **C.** *cysl-1* is required for Suh phenotype of *skn-1(uwa02gf)*. Animals of each genotype were exposed to 50 ppm H_2_S overnight and the number of animals that survived was scored. Data are from 6-13 independent experiments with at least 10 animals in each cohort. Error bars are SEM. **D.** *cysl-1* is necessary for expression of RHY-1 to rescue *hif-1* lethality in H_2_S. Extrachromosomal arrays of *rhy-1* were crossed into each strain. The viability of animals carrying the array (filled bars) and siblings that had lost the array (open bars) was scored after exposure to 50 ppm H_2_S overnight. **E.** Expression of CYSL-1 cannot rescue *hif-1* lethality in H_2_S. Animals were incubated in 50 ppm H_2_S overnight, and then the number of surviving animals was scored. Ex[CYSL-1] are transgenic animals that contain an extrachromosomal array with *cysl-1* genomic DNA.

Stabilization of HIF-1 by loss-of-function mutations in *rhy-1* increases expression of *hif-1*-target genes and disrupts a behavioral response in which worms increase locomotion when moved to room air after a short exposure to anoxia (Ma et al., 2012; Shen et al., 2006). These *rhy-1* phenotypes are suppressed by mutations in *cysl-1*, a member of the cystathionine-beta synthase/cysteine synthase family of type-II pyridoxal-5’-phosphate (PLP) dependent proteins with O-acetylserine sulfhydrase activity *in vitro* (Vozdek et al., 2013). In H_2_S, CYSL-1 has been shown to physically interact with the EGL-9 prolyl hydroxylase to prevent it from hydroxylating HIF-1, thereby leading to the stabilization of HIF-1 protein in H_2_S (Ma et al., 2012). We therefore considered the possibility that *rhy-1* also acts with *cysl-1* in the *hif-1*-independent response to H_2_S.

CYSL-1 physically interacts with EGL-9 and this interaction is stimulated by H_2_S and negatively regulated by RHY-1 (Figure 5A, Ma et al., 2012). EGL-9 is a negative regulator of HIF-1, but there is also evidence that EGL-9 has roles independent of its HIF-1 regulatory activity (Shao et al., 2009). We therefore tested whether EGL-9 worked with RHY-1 and CYSL-1 independently of *hif-1* in H_2_S. We found that activation of SKN-1 by RNAi of *wdr-23* could rescue *hif-1(ia04); egl-9 (sa307)* double-mutant animals exposed to H_2_S (Figure 5B). This result shows that *egl-9* is not required for increased SKN-1 activity to bypass the requirement for *hif-1* in H_2_S, implying that the physical interaction between CYSL-1 and EGL-9 is not involved.

We next asked whether *cysl-1* is required, like *rhy-1*, for the SKN-1-mediated Suh phenotype. As has been previously reported, we found that *cysl-1(ok762)* mutant animals die when exposed to 50 ppm H_2_S (Budde and Roth, 2011). This lethality was suppressed by RNAi of *egl-9* (Figure 5E), consistent with *cysl-1* acting to positively regulate *hif-1* upstream of *egl-9* (Ma et al., 2012). In contrast, activation of SKN-1 by *wdr-23* RNAi did not rescue *cysl-1(ok762)* mutant animals exposed to H_2_S (Figure 5B). Therefore, both *rhy-1* and *cysl-1* are required for the Suh phenotype in animals with increased activation of SKN-1. Corroborating this observation, *cysl-1(ok762); hif-1(ia04); skn-1(uwa02)* triple mutants died when exposed to H_2_S (Figure 5C). We conclude that *cysl-1* is necessary for the RHY-1 to promote survival in hif-1(ia04) mutant animals exposed to H_2_S.

In order to determine if expression of CYSL-1 was sufficient to bypass the requirement for *hif-1* in H_2_S we generated a *cysl-1* overexpression transgenic line by amplifying genomic *cysl-1* with ~2 KB upstream to include the native promoter region and ~500 bp downstream of the genomic locus (Wormbase WS204, http://ws204.wormbase.org/). This construct produced functional CYSL-1, as it rescued the H_2_S-sensitivity of *cysl-1(ok762)* mutant animals (Figure 5E). In contrast to overexpression of RHY-1, CYSL-1 overexpression did not rescue *hif-1(ia04)* lethality in H_2_S (figure 5E). We conclude that *cysl-1* is necessary but not sufficient for SKN-1 activity to bypass the requirement of *hif-1* in H_2_S.

## Discussion

We have found that expression of RHY-1 can bypass the requirement for HIF-1 in H_2_S. This is remarkable, since *hif-1* is required for all observed initial changes in gene expression induced by a short exposure to H_2_S (Miller et al., 2011). The conserved HIF-1 transcription factor has been studied in depth in a variety of organisms; however, to our knowledge, there have not been other genetic factors identified that can suppress the effects of *hif-1* loss in any context.

The mutations we recovered do not rescue *hif-1* phenotypes in hypoxia or cyanide, suggesting genetically independent roles for HIF-1 in different stress conditions. This is consistent with our previous observation that there was little overlap in *hif-1*-dependent changes in gene expression when *C. elegans* are exposed to hypoxia or H_2_S (Miller et al., 2011). However, these results also suggest that toxicity of H_2_S and HCN result from fundamentally different mechanisms, as RHY-1 can promote survival in H_2_S but not HCN. It is generally posited that H_2_S and HCN toxicity arise from inhibition of cytochrome c oxidase, though the mechanisms underlying cytotoxicity are not fully understood (Jiang et al., 2016). Indeed, it has long been appreciated that there are distinct requirements for treatment of H_2_S and HCN poisoning (Smith and Gosselin, 1976).

One mechanism by which HIF-1 promotes survival in H_2_S is by induction of SQRD-1, a sulfide-quinone oxidoreductase that can oxidize H_2_S (Budde and Roth, 2011; Shibata et al., 1999). The SQRD-1 sulfide-quinone oxidoreductase activity is conserved from bacteria to mammals (Ackermann et al., 2011). Oxidation of H_2_S by sulfide-quinone oxidoreductase is the first step in metabolism of cellular H_2_S, followed by further oxidation by the ETHE-1 sulfur dioxygenase (Quinzii et al., 2017). Decreased activity of ETHE-1 causes ethylmalonic encephalopathy resulting from accumulation of endogenously produced H_2_S in mammals (Tiranti et al., 2009), and *C. elegans* with mutations in either *sqrd-1* or *ethe-1* die when exposed to low H_2_S (Budde and Roth, 2011). We have found that RHY-1 can suppress lethality of *sqrd-1* mutant animals. This result suggests that there may be a novel H_2_S metabolism pathway that does not require oxidation by sulfide-quinone oxidoreductase.

Although there are a variety of acyl-transferases with unknown functions coded for in the human genome, the closest orthologue to RHY-1, ACYL3, has been lost in the primate lineage after the divergence of gorillas from the human lineage (Zhu et al., 2007). In wild-type animals exposed to H_2_S, RHY-1 negatively regulates HIF-1 by preventing CYSL-1 from inhibiting EGL-9 (Ma et al., 2012). In contrast, we have found that RHY-1 and CYSL-1 are both necessary to promote survival in H_2_S independently of HIF-1. It is not clear how RHY-1, an integral membrane protein in the endoplasmic reticulum (Shen et al., 2006), regulates the activity of CYSL-1, a cytoplasmic enzyme. Understanding the mechanistic basis of this RHY-1 activity could lead to the development of novel treatments for H_2_S poisoning and could provide a therapeutic target to modulate the activity of H_2_S signaling.

## Materials and Methods

### Strains

*Caenorhabditis elegans* strains were cultured at 20 °C on NGM plates with OP50 *E. coli*. Strains used were: ZG31 [*hif-1(ia04)* V], RB1297 [*rhy-1(ok1402)* II], RB899 [*cysl-1(ok762)* X], CB6088 [*egl-9(sa307); hif-1(ia4) V*.], CL2166 *(dvIs19 [(pAF15)gst-4p::GFP::NLS]* III), and *sqrd-1(tm3378)* V. Strains SPC167 [*dvIs19 III; skn-1(lax120) IV]* and SPC168 [*dvIs19 III; skn-1(lax188) IV]* were provided by Dr. Sean Curran (University of Southern California). Strains generated in this work are *hif-1(ia04); skn-1(uwa02*) IV, DLM23 *hif-1(ia04); skn-1(uwa06)* IV, DLM22 *hif-1(ia04); wdr-23(uwa05)* I, *hif-1(ia04); wdr-23(uwa13)* I, and DLM24 *hif-1(ia04); wdr-23(uwa15)* I. Double and triple mutants were generated using standard genetic techniques, and genotypes were verified by PCR genotyping. Primer sequences are available upon request.

Transgenic strains were generated as in (Mello et al., 1991). To generate extrachromosomal arrays, 10ng/µl of the gene of interest, amplified by PCR from genomic DNA, and co-injection reporter (Pmyo-2::RFP or Pmyo-3::RFP) were co-injected with Yeast Centromere Plasmid prs415 filler DNA to a final concentration of 100 ng/µl. Genomic overexpression constructs were amplified from N2 genomic DNA with primers to include 1-2 kb upstream and downstream of coding regions.

### H_2_S exposure

*C. elegans* were exposed to H_2_S in atmospheric boxes perfused with H_2_S continuously diluted into room air to generate stable atmospheres with 50 or 150 ppm H_2_S in room air, as previously described (Cox and Spradling, 2006). 50 ppm H_2_S was used in all experiments unless otherwise stated. Source tanks of compressed H_2_S gas (5,000 ppm balanced with N_2_) were purchased from Airgas (Seattle, WA). Mixing was achieved using SmartTrak mass flow controllers (Sierra Instruments). H_2_S concentrations were verified with PowerLab (ADinstruments) detection of H_2_S as compared to a verified 100 ppm H_2_S standard (Airgas) as described (Miller and Roth, 2007). Experiments were conducted at room temperature. Matched controls were perfused with room air and maintained at the same temperature in the hood. To score survival, 10-20 L4 animals were picked to NGM plates with OP50 and exposed to 50ppm H_2_S for 16 hours. Survival was scored immediately after removal from the H_2_S atmosphere.

### EMS mutagenesis

*hif-1(ia04) C. elegans* were synchronized by bleaching and grown to L4. 20 µL of EMS was added to 4mL of worms in M9 buffer and rotated for 4 hours at room temperature. The mutagenized worms were pelleted and washed with M9 with 0.01% SDS to prevent worms from sticking to the tube and plated on HG plates to recover overnight. The gravid mutagenized P0 animals were bleached and the F1 progeny were grown on HG plates to adulthood, then bleached to isolate F2 embryos. F2 eggs were plated on NGM plates with OP50 and grow to L4 in room air. The F2 L4s were exposed to 50ppm H_2_S for 16 hours. Animals that survived H_2_S exposure were singled and re-tested for survival 3 times. These surviving animals were then outcrossed 4 or more times before use.

### Whole Genome Sequencing and Analysis

Genomic DNA was prepared with Puregene Core Kit A (Qiagen). When necessary, DNA was further purified by phenol-chloroform extraction and ethanol precipitation. Samples were sequenced at the University of Utah Sequencing Core and analyzed using a modified Cloudmap workflow on usegalaxy.org to generate a list of SNPs as compared to the parent *hif-1(ia04)* strain (Minevich et al., 2012).

### Quantitative RT-PCR

Total RNA was isolated from ~9000 bleach-synchronized young adult *C. elegans*. For H_2_S exposed samples, young adult *C. elegans* were exposed to 50 ppm H_2_S for 1 hour prior to RNA harvest. Alternatively, mixed populations of *C. elegans* were harvested off 10 cm NGM plates for *rhy-1* qRT-PCR. Animals were harvested in M9 buffer, and 100µL of packed animals added to 1mL TRIzol RNA isolation reagent (Life Technnologies), and flash frozen in liquid nitrogen. mRNA was isolated following the manufacturer’s protocol, and then cDNA was synthesized from 5µg RNA using polyT primers included with Superscript III Reverse Transcriptase (Invitrogen) according to the manufacturer’s instructions. Each 10µL qPCR reaction contained 1µL cDNA and 5µL 2X Sybr Green Master Mix (Kappa Biosystems). Primers were added using a 0.2µL pin tool. Absorbance was measured over 40 cycles using a Mastercycler RealPlex 2 (Eppendorf). The threshold cycle (C_t_) for each sample was measured using the provided software, and normalized to *hil-1* and *irs-2* controls to generate ΔC_t_ values as previously described (Miller et al., 2011). ΔΔC_t_ was calculated as the change in ΔC_t_ between animals of the same genotype exposed to H_2_S and room air controls or between *C. elegans* strains exposed to the same conditions. Figures show average ΔΔC_t_ ± standard deviation for at least 3 independent replicates.

### H_2_S survival

For survival assays, a minimum of 10 L4 *C. elegans* were picked to a 3 cm NGM plate seeded with OP50 bacteria. The plates were then exposed to 50 ppm H_2_S for 16 hours and survival was scored by visual inspection immediately after removal from H_2_S. Animals were considered dead if they did not respond to prodding with a platinum wire. For RNAi H_2_S survival, 3 cm RNAi plates were seeded with 30µL of log-phase RNAi bacterial culture. The next day, gravid animals were placed in a spot of bleach on the RNAi plates and allowed to grow at 20 °C until L4. The L4 animals were then exposed to H_2_S for 16 hours and survival was scored upon removal from H_2_S.

### Hypoxia response assays

Hypoxic environments were constructed by continuously flowing 5,000 ppm O_2_ (balance N_2_) through environmental boxes, as previously described (Fawcett et al., 2012). Premixed tanks with 5,000 ppm O_2_/N_2_ were purchased from Airgas (Seattle, WA) and certified standard to be within 2% of the desired O_2_ concentration.

To measure the rate of egg-laying in hypoxia, 3 cm plates with OP50 were ringed with palmitic acid to create a physical barrier to keep *C. elegans* on the agar media as in (Miller and Roth, 2009). 1-6 adult worms were picked to each plate and exposed to 5000 ppm O_2_ for 20 h and number of eggs laid per was scored. Plates in which all the adult animals could not be accounted for were censored and excluded from the analysis.

To assay embryo viability in hypoxia, gravid adult worms were chopped with a razor blade in a drop of M9 buffer on a glass slide. The dissected embryos were transferred to a 3 cm NGM plate by mouth pipet and then exposed to hypoxia for 20 h. After hypoxic exposure, the number of eggs that had not hatched and the number of larvae were counted to calculate viability.

### Cyanide viability assay

To generate gaseous environments with defined concentrations of HCN, 125 mg of NaCN was freshly dissolved in 10mL M9. In a fume hood, defined amounts of cyanide were placed into the empty lid of a 3cm plate in a 2.5L Anaeropack box (Mitsubishi Gas Chemical Co., Inc.) that contained plates with 10-20 L4 animals of each strain on OP50. An equal volume of 14 M HCl was added to the drop of NaCN to volatilize HCN, and the box was rapidly sealed. The amount of HCN generated was calculated assuming 100% conversion of NaCN to gaseous HCN in the 2.5 L box. After 16 h, the box was opened and plates removed to score the number of animals that survived.

### *Pgst-4*::GFP expression

L4 *C. elegans* were picked into 50 mM sodium azide to paralyze the animals and placed on a 2% agarose pad for visualization. *C. elegans* were visualized on a Nikon Eclipse 90i and pictures were taken with an Andor Zyla sCMOS camera.

## Acknowledgements

We thank members of the Miller Lab for fruitful discussions and critical reading of the manuscript. This work was funded by R00 AG033047 (to DLM) and R01 ES024958 (to DLM) and the Glenn Foundation for Biomedical Research. JH was supported by the Genetics Approaches to Aging Training Grant NIH/NIA T32 AG000057. Some strains were provided by the CGC, which is funded by NIH Office of Research Infrastructure Programs (P40 OD010440). DLM is a New Scholar in Aging of the Ellison Medical Foundation.

---

**Supplemental Figure 1.**
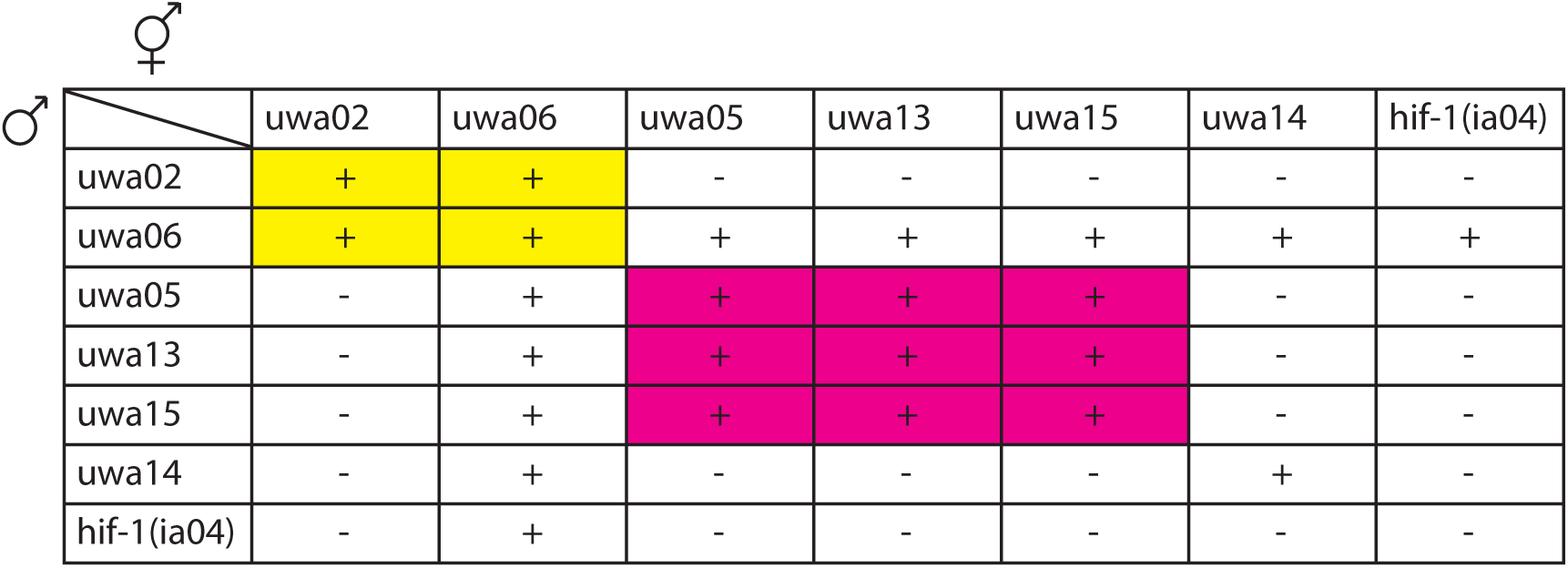
Complementation analysis of *suh* alleles. Progeny of matings between males (as indicated on the left column) and hermaphrodites with the genotype indicated in the top row were exposed to H_2_S. If a majority of heterozygous animals survived H_2_S, indicating non-complementation, the cells are marked with +. Complementing crosses are indicated with –. The two independent complementation groups are colored in cyan and magenta.

**Supplemental Figure 2.**
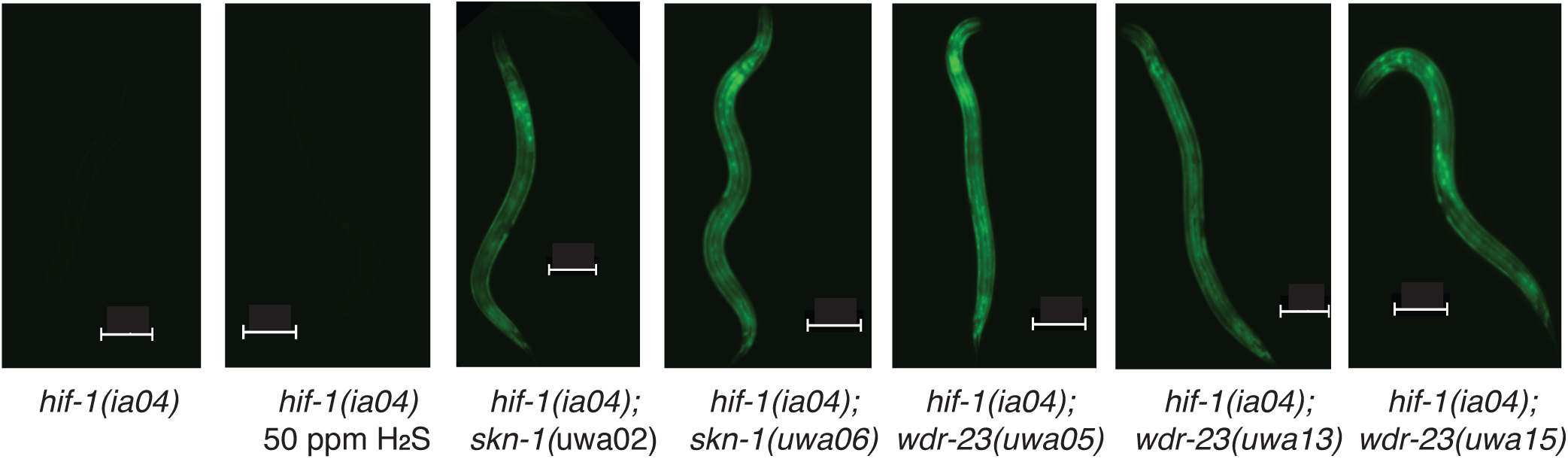
*suh* mutations increase SKN-1 activity. Representative images of animals with the *Pgst-4::gfp* transgene. There was no expression of GFP in *hif-1(ia04)* animals, even after exposure to 50 ppm H_2_S. Sscale bar is 100 µM.

**Supplemental Figure 3.**
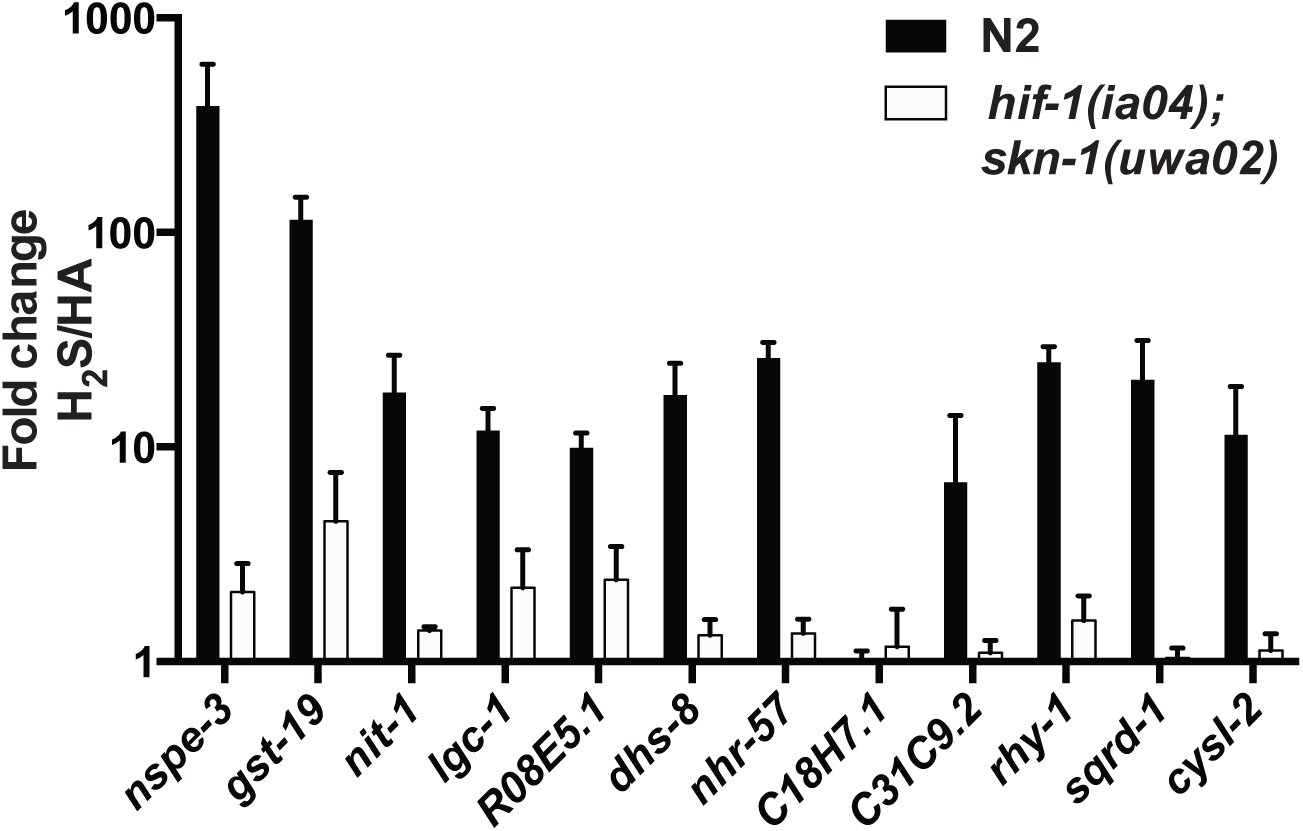
H_2_S-induced gene expression is not restored in *skn-1(uwa02gf)* mutant animals. Change in transcript abundance of H_2_S-inducible gene products measured by qRT-PCR after exposure to 50 ppm H_2_S for 1 h. Average fold change calculated from ΔΔC_t_ (ΔC_t_^H2S^-ΔC_t_^RA^), error bars show standard deviation. N=3 independent experiments each with three technical replicates for both N2 and *hif-1(ia04); skn-1(uwa02)*.

**Supplemental Figure 4.**
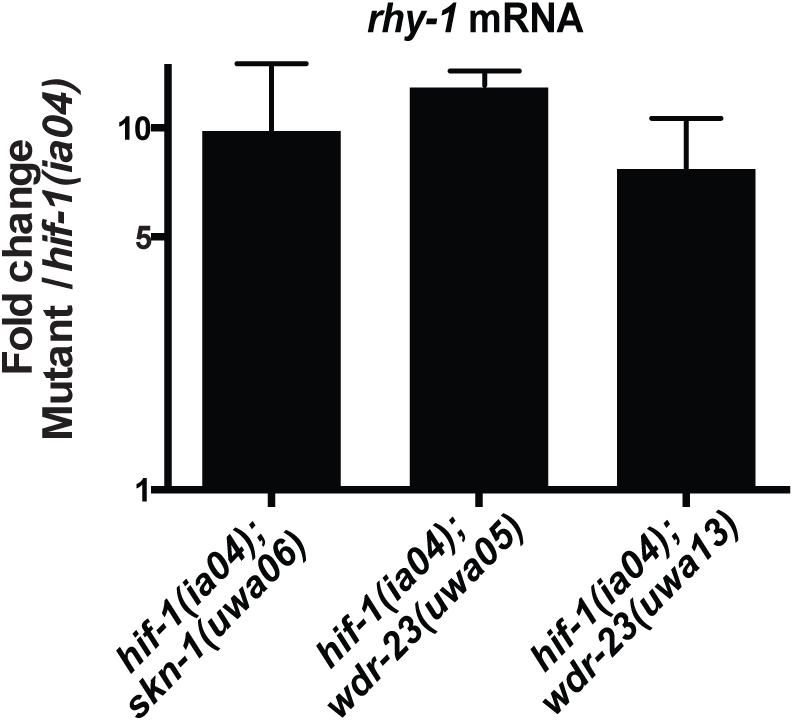
Increased experession of *rhy-1* in *suh* mutants. *rhy-1* mRNA abundance in *suh* mutant animals, as measured by qRT-PCR. Average fold change in *rhy-1* expression between the indicated Suh strain compared to *hif-1(ia04)*. Bars indicated one standard deviation.

## References

Ackermann, M., M. Kubitza, K. Maier, A. Brawanski, G. Hauska, and A. L. Piña. 2011. The vertebrate homolog of sulfide-quinone reductase is expressed in mitochondria of neuronal tissues. Neuroscience. 199:1–12.

An, J. H., and T. K. Blackwell. 2003. SKN-1 links *C. elegans* mesendodermal specification to a conserved oxidative stress response. Genes Dev. 17:1882–1893.

An, J. H., K. Vranas, M. Lucke, H. Inoue, N. Hisamoto, K. Matsumoto, and T. K. Blackwell. 2005. Regulation of the *Caenorhabditis elegans* oxidative stress defense protein SKN-1 by glycogen synthase kinase-3. PNAS. 102:16275–16280.

Blackstone, E., and M. B. Roth. 2007. Suspended animation-like state protects mice from lethal hypoxia. Shock. 27:370–372.

Bowerman, B., B. A. Eaton, and J. R. Priess. 1992. *skn-1*, a maternally expressed gene required to specify the fate of ventral blastomeres in the early *C. elegans* embryo. Cell. 68:1061–1075.

Budde, M. W., and M. B. Roth. 2011. The response *of* Caenorhabditis elegans to hydrogen sulfide and hydrogen cyanide. Genetics. 189:521–532.

Budde, M. W., and M. B. Roth. 2010. Hydrogen Sulfide Increases Hypoxia-inducible Factor-1 Activity Independently of von Hippel, ÄìLindau Tumor Suppressor-1 in *C. elegans*. Mol Biol Cell. 21:212–217.

Calvert, J. W., S. Jha, S. Gundewar, J. W. Elrod, A. Ramachandran, C. B. Pattillo, C. G. Kevil, and D. J. Lefer. 2009. Hydrogen sulfide mediates cardioprotection through Nrf2 signaling. Circ Res. 105:365–374.

Choe, K. P., A. J. Przybysz, and K. Strange. 2009. The WD40 repeat protein WDR-23 functions with the CUL4/DDB1 ubiquitin ligase to regulate nuclear abundance and activity of SKN-1 in *Caenorhabditis elegans*. Mol Cell Biol. 29:2704–2715.

Cooper, C. E., and G. C. Brown. 2008. The inhibition of mitochondrial cytochrome oxidase by the gases carbon monoxide, nitric oxide, hydrogen cyanide and hydrogen sulfide: chemical mechanism and physiological significance. J Bioenerg Biomembr. 40:533–539.

Cox, R. T., and A. C. Spradling. 2006. Milton controls the early acquisition of mitochondria by *Drosophila* oocytes. Development. 133:3371–3377.

Darby, C., C. L. Cosma, J. H. Thomas, and C. Manoil. 1999. Lethal paralysis of *Caenorhabditis elegans* by *Pseudomonas aeruginosa*. PNAS. 96:15202–15207.

Epstein, A. C., J. M. Gleadle, L. A. McNeill, K. S. Hewitson, J. O’Rourke, D. R. Mole, M. Mukherji, E. Metzen, M. I. Wilson, A. Dhanda, Y. M. Tian, N. Masson, D. L. Hamilton, P. Jaakkola, R. Barstead, J. Hodgkin, P. H. Maxwell, C. W. Pugh, C. J. Schofield, and P. J. Ratcliffe. 2001. *C. elegans* EGL-9 and mammalian homologs define a family of dioxygenases that regulate HIF by prolyl hydroxylation. Cell. 107:43–54.

Fawcett, E. M., J. M. Hoyt, J. K. Johnson, and D. L. Miller. 2015. Hypoxia disrupts proteostasis in *Caenorhabditis elegans*. Aging Cell. 14:92–101.

Fawcett, E. M., J. W. Horsman, and D. L. Miller. 2012. Creating Defined Gaseous Environments to Study the Effects of Hypoxia on C. elegans. J Vis Exp. (65), e4088 10.3791/4088.

Gallagher, L. A., and C. Manoil. 2001. *Pseudomonas aeruginosa* PAO1 kills *Caenorhabditis elegans* by cyanide poisoning. J Bacteriol. 183:6207–6214.

Hasegawa, K., and J. Miwa. 2010. Genetic and cellular characterization of *Caenorhabditis elegans* mutants abnormal in the regulation of many phase II enzymes. PLoS One. 5:e11194.

Hildebrandt, T. M., and M. K. Grieshaber. 2008. Three enzymatic activities catalyze the oxidation of sulfide to thiosulfate in mammalian and invertebrate mitochondria. FEBS J. 275:3352–3361.

Horsman, J. W., and D. L. Miller. 2015. Mitochondrial sulfide quinone oxidoreductase prevents activation of the unfolded protein response in hydrogen sulfide. J. Biol. Chem. 291:5320–5325.

Jensen, A. R., N. A. Drucker, S. Khaneki, M. J. Ferkowicz, M. C. Yoder, E. R. DeLeon, K. R. Olson, and T. A. Markel. 2017. Hydrogen Sulfide: A Potential Novel Therapy for the Treatment of Ischemia. Shock. 48:511–524.

Jiang, H., R. Guo, and J. A. Powell-Coffman. 2001. The *Caenorhabditis elegans hif-1* gene encodes a bHLH-PAS protein that is required for adaptation to hypoxia. PNAS. 98:7916–7921.

Jiang, J., A. Chan, S. Ali, A. Saha, K. J. Haushalter, W. L. Lam, M. Glasheen, J. Parker, M. Brenner, S. Mahon, H. H. Patel, R. Ambasudhan, S. A. Lipton, R. B. Pilz, and G. R. Boss. 2016. Hydrogen Sulfide--Mechanisms of Toxicity and Development of an Antidote. Sci Rep. 6:20831.

Kell, A., N. Ventura, N. Kahn, and T. E. Johnson. 2007. Activation of SKN-1 by novel kinases in *Caenorhabditis elegans*. Free Rad Biol Med. 43:1560–1566.

Kobayashi, A., M. I. Kang, H. Okawa, M. Ohtsuji, Y. Zenke, T. Chiba, K. Igarashi, and M. Yamamoto. 2004. Oxidative stress sensor Keap1 functions as an adaptor for Cul3-based E3 ligase to regulate proteasomal degradation of Nrf2. Mol Cell Biol. 24:7130–7139.

Li, L., P. Rose, and P. K. Moore. 2011. Hydrogen sulfide and cell signaling. Annu Rev Pharmacol Toxicol. 51:169–187.

Ma, D. K., R. Vozdek, N. Bhatla, and H. R. Horvitz. 2012. CYSL-1 interacts with the O2-sensing hydroxylase EGL-9 to promote H2S-modulated hypoxia-induced behavioral plasticity in *C. elegans*. Neuron. 73:925–940.

Malone Rubright, S. L., L. L. Pearce, and J. Peterson. 2017. Environmental toxicology of hydrogen sulfide. Nitric Oxide. 71:1–13.

Maxwell, P. H., M. S. Wiesener, G. W. Chang, S. C. Clifford, E. C. Vaux, M. E. Cockman, C. C. Wykoff, W. Pugh, E. R. Maher, and P. J. Ratcliffe. 1999. The tumour suppressor protein VHL targets hypoxia-inducible factors for oxygen-dependent proteolysis. Nature. 399:271–275.

Mello, C. C., J. M. Kramer, D. Stinchcomb, and V. Ambros. 1991. Efficient gene transfer in *C.elegans*: extrachromosomal maintenance and integration of transforming sequences. EMBO J. 10:3959–3970.

Miller, D. L., M. W. Budde, and M. B. Roth. 2011. HIF-1 and SKN-1 coordinate the transcriptional response to hydrogen sulfide in *Caenorhabditis elegans*. PLoS One. 6:e25476.

Miller, D. L., and M. B. Roth. 2009. *C. elegans* Are Protected from Lethal Hypoxia by an Embryonic Diapause. Curr Biol. 19:1233–1237.

Miller, D. L., and M. B. Roth. 2007. Hydrogen sulfide increases thermotolerance and lifespan in *Caenorhabditis elegans*. PNAS. 104:20618–20622.

Minevich, G., D. S. Park, D. Blankenberg, R. J. Poole, and O. Hobert. 2012. CloudMap: a cloud-based pipeline for analysis of mutant genome sequences. Genetics. 192:1249–1269.

Nystul, T. G., and M. B. Roth. 2004. Carbon monoxide-induced suspended animation protects against hypoxic damage in *Caenorhabditis elegans*. PNAS. 101:9133–9136.

Paek, J., J. Y. Lo, S. D. Narasimhan, T. N. Nguyen, K. Glover-Cutter, S. Robida-Stubbs, T. Suzuki, M. Yamamoto, T. K. Blackwell, and S. P. Curran. 2012. Mitochondrial SKN-1/Nrf mediates a conserved starvation response. Cell Metab. 16:526–537.

Quinzii, C. M., M. Luna-Sanchez, M. Ziosi, A. Hidalgo-Gutierrez, G. Kleiner, and L. C. Lopez. 2017. The Role of Sulfide Oxidation Impairment in the Pathogenesis of Primary CoQ Deficiency. Frontiers in Physiology. 8:525.

Semenza, G. L. 2012. Hypoxia-inducible factors in physiology and medicine. Cell. 148:399–408.

Shao, Z., Y. Zhang, and J. A. Powell-Coffman. 2009. Two distinct roles for EGL-9 in the regulation of HIF-1-mediated gene expression in *Caenorhabditis elegans*. Genetics. 183:821–829.

Shao, Z., Y. Zhang, Q. Ye, J. N. Saldanha, and J. A. Powell-Coffman. 2010. *C. elegans* SWAN-1 Binds to EGL-9 and Regulates HIF-1-Mediated Resistance to the Bacterial Pathogen *Pseudomonas aeruginosa* PAO1. PLoS Pathog. 6(8): e1001075. doi:10.1371/journal.ppat.1001075.

Shen, C., Z. Shao, and J. A. Powell-Coffman. 2006. The Caenorhabditis elegans *rhy-1* gene inhibits HIF-1 hypoxia-inducible factor activity in a negative feedback loop that does not include *vhl-1*. Genetics. 174:1205–1214.

Shen, C., and J. A. Powell-Coffman. 2003. Genetic Analysis of Hypoxia Signaling and Response in *C. elegans*. Ann NY Acad Sci. 995:191–199.

Shibata, H., M. Takahashi, I. Yamaguchi, and S. Kobayashi. 1999. Sulfide oxidation by gene expressions of sulfide-quinone oxidoreductase and ubiquinone-8 biosynthase in *Escherichia coli*. J Biosci Bioeng. 88:244–249.

Smith, R. P., and R. E. Gosselin. 1976. Current concepts about the treatment of selected poisonings: nitrite, cyanide, sulfide, barium, and quinidine. Annu Rev Pharmacol Toxicol. 16:189–199.

Tang, L., and K. P. Choe. 2015. Characterization of *skn-1*/*wdr-23* phenotypes in *Caenorhabditis elegans*; pleiotrophy, aging, glutathione, and interactions with other longevity pathways. Mech Ageing Dev. 149:88–98.

Tiranti, V., C. Viscomi, T. Hildebrandt, I. Di Meo, R. Mineri, C. Tiveron, M. D. Levitt, A. Prelle, G. Fagiolari, M. Rimoldi, and M. Zeviani. 2009. Loss of ETHE1, a mitochondrial dioxygenase, causes fatal sulfide toxicity in ethylmalonic encephalopathy. Nat Med. 15:200–205.

Vozdek, R., A. Hnízda, J. Krijt, L. Será, and V. Kožich. 2013. Biochemical properties of nematode O-acetylserine(thiol)lyase paralogs imply their distinct roles in hydrogen sulfide homeostasis. Biochim Biophys Acta. 1834:2691–2701.

Walker, A. K., R. See, C. Batchelder, T. Kophengnavong, J. T. Gronniger, Y. Shi, and T. K. Blackwell. 2000. A conserved transcription motif suggesting functional parallels between *Caenorhabditis elegans* SKN-1 and Cap’n’Collar-related basic leucine zipper proteins. J Biol Chem. 275:22166–22171.

Wu, D., J. Wang, H. Li, M. Xue, A. Ji, and Y. Li. 2015. Role of Hydrogen Sulfide in Ischemia-Reperfusion Injury. Oxid Med Cell Longev. 2015:186908.

Yang, G., K. Zhao, Y. Ju, S. Mani, Q. Cao, S. Puukila, N. Khaper, L. Wu, and R. Wang. 2013. Hydrogen sulfide protects against cellular senescence via S-sulfhydration of Keap1 and activation of Nrf2. Antioxid Redox Signal. 18:1906–1919.

Zhu, J., J. Z. Sanborn, M. Diekhans, C. B. Lowe, T. H. Pringle, and D. Haussler. 2007. Comparative genomics search for losses of long-established genes on the human lineage. PLoS Comput Biol. 3:e247.

